# Phenome-wide search for pleiotropic loci highlights key genes and molecular pathways for human complex traits

**DOI:** 10.1101/672758

**Authors:** Anton E. Shikov, Alexander V. Predeus, Yury A. Barbitoff

## Abstract

Over recent decades, genome-wide association studies (GWAS) have dramatically changed the understanding of human genetics. A recent genetic data release by UK Biobank has allowed many researchers worldwide to have comprehensive look into the genetic architecture of thousands of human phenotypes. In this study, we developed a novel statistical framework to assess phenome-wide significance and genetic pleiotropy across the human phenome based on GWAS summary statistics. We demonstrate widespread sharing of genetic architecture components between distinct groups of traits. Apart from known multiple associations inside the MHC locus, we discover high degree of pleiotropy for genes involved in immune system function, apoptosis, hemostasis cascades, as well as lipid and xenobiotic metabolism. We find several notable examples of novel pleiotropic loci (e.g., the *MIR2113* microRNA broadly associated with cognition), and provide several possible mechanisms for these association signals. Our results allow for a functional phenome-wide look into the shared components of genetic architecture of human complex traits, and highlight crucial genes and pathways for their development.

## Introduction

Over the recent decades, many advances have been made in uncovering the genetic architecture of human complex traits. Genome-wide association studies (GWAS), emerging at the onset of the millennium, have facilitated the discovery of susceptibility loci for both complex disease and quantitative traits (reviewed in Mills and Rahal^1^). Most recent GWAS for common phenotypes such as type 2 diabetes include up to 900,000 individuals and allow to identify numerous novel risk loci^2^. Many methods have been developed to interpret the GWAS signal on both single-variant and genome scale, including a group of specialized gene set enrichment tests^3–5^ aimed at discovery of biological pathways and mechanisms related to complex traits. Integration of GWAS and expression quantitative trait loci (eQTL) data has also shed light onto the role of gene expression in complex traits, and allowed for identification and validation of causal genes^6^.

Advances in biobanking, genetic data acquisition, and analysis have allowed first glimpse into the genetic architecture of multiple traits at once, with the UK Biobank (UKB) genetic dataset of 500,000 individuals being extensively used for diverse studies in statistical genetics^7^. These included, for example, estimation of natural selection effects on complex trait-associated variation^8^, and development of trait-specific risk scores for identification of individuals at high risk for complex disease^9^. Genetic associations in the UK Biobank data have attracted the attention of researchers worldwide, and several efforts have been made to aggregate genetic association data across all traits^10–12^. These studies highlighted the complexity of the human phenome, and pinpointed notable loci with multiple associations (e.g., the MHC locus).

Several attempts have been made to assess the association of individual variants or genes across the whole phenome (phenome-wide association studies, PheWAS) (e.g., Schmidt *et al*^13^), though no unified statistical framework to interpret such phenome-wide associations has been proposed. Pleiotropy of trait-associated variants in the human genome has also attracted lots of attention in the field; and Mendelian randomization based approaches have been proposed to detect pleiotropy in GWAS data^14^. At the same time, UK Biobank dataset offers an excellent opportunity to analyze shared components of the genetic architecture of human phenotypes on the phenome-scale, with global overviews suggesting widespread signals of pleiotropy in the UK Biobank data^15,16^. In this study, we developed a statistical framework to explore the landscape of phenome-wide associations in GWAS summary statistics derived from UK Biobank dataset, and identified multiple shared blocks of genetic architecture of diverse human complex traits.

## Results

### Complexity reduction by clustering of similar traits

To conduct a genome-wide scan for pleiotropic loci, we first obtained sets of significantly associated SNPs for all phenotypes in the UK Biobank data using pre-calculated GWAS summary statistics provided by Benjamin Neale’s lab (release 1, downloaded 2018-02-25). The dataset included both standard GWAS loci as well as imputed variants, totaling 10,894,597 variants. We focused our analysis only on 543 complex traits that have significant non-zero partitioned heritability estimates (h^2^). A total of 469,013 (4.27%) SNPs had at least one phenotype associated at genome-wide significant level, with an average of 4.34 phenotypes associated with each locus. Interestingly, we observed numerous multiple associations across the dataset, with 230296 (49.21%) of SNPs having > 1 associated phenotype, and more than 10 associated phenotypes for 57,856 (12.34%) SNPs. While many of these multiple associations could be true pleiotropic variants, much of the signal likely arises from statistical artifacts or multiple highly correlated phenotypes. Hence, we went on to cluster traits that share a significant proportion of their genetic architecture. We applied the Jaccard distance metric as a simple and efficient way that can group traits based on pre-selected significant SNPs. Hierarchical clustering based on Jaccard distance between phenotypes identified 307 independent trait clusters with 1.76 phenotypes on average. Among these, 29 clusters comprised 3 or more phenotypes (see Extended Data Figure 1). The clustering procedure has dramatically decreased the amount of SNPs having multiple associations (92,224 (19.7%) SNPs with more than one association compared to 230,296 in non-clustered data) (see Extended Data Figure 2); as well as the average number of associated trait clusters (1.33 cluster per variant). Notably the number of pleiotropic SNPs with more than 10 associations has dramatically dropped to 0.014% (64 variants) after performing clusterization procedure (Figure 1d-e). Interestingly, variants that have > 5 associated clusters also have higher minor allele frequencies, i.e. most rare variants are not pleiotropic (Figure 1e). Similar results were obtained when comparing SNPs with less or more than 15 associated individual phenotypes (see Extended Data Figure 3), indicating that the clustering procedure didn’t affect the relationship between MAF and pleiotropy.

**Figure 1.**
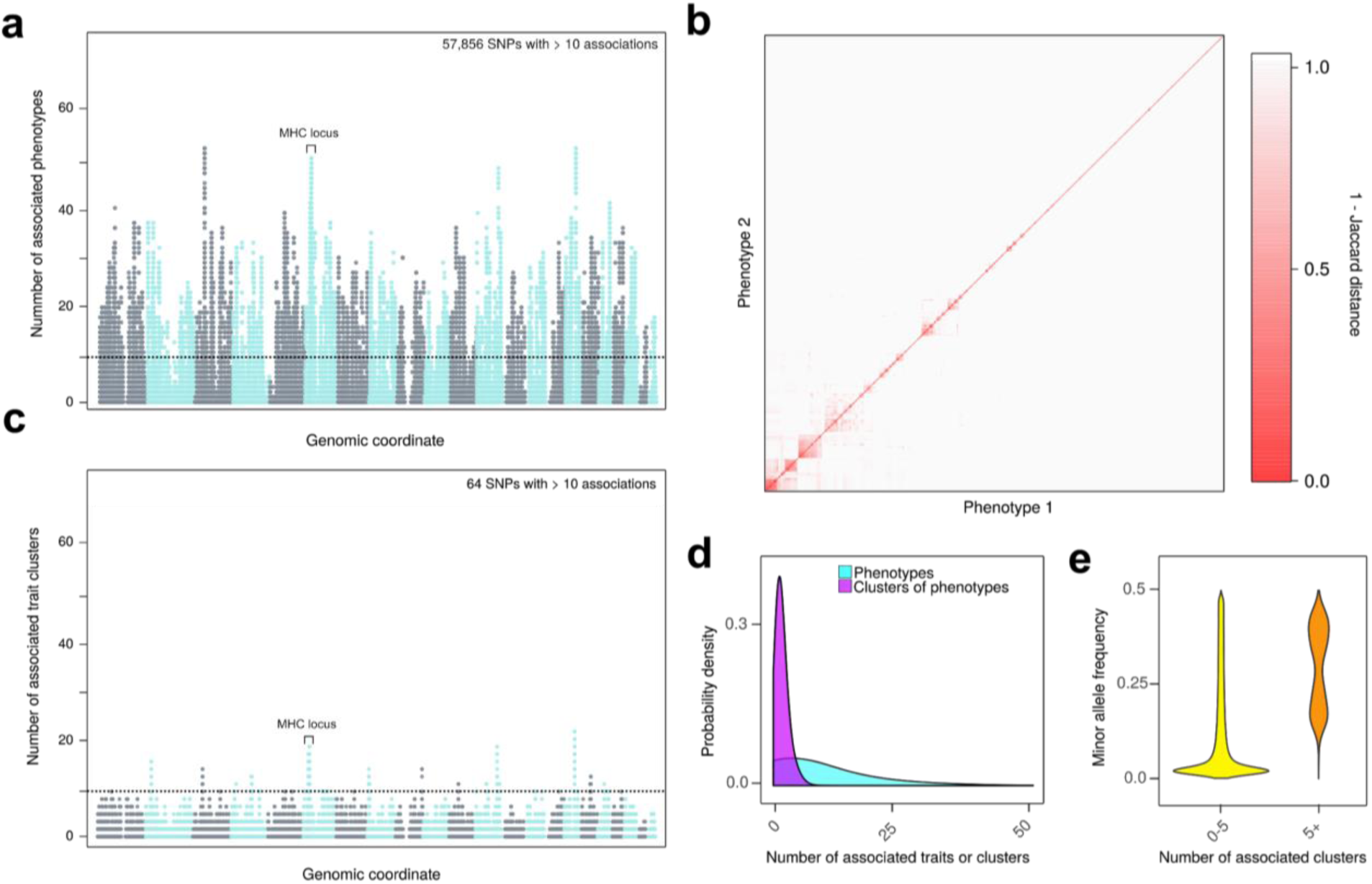
Clustering of similar traits significantly reduces the number of multiple associations. **a** Manhattan plot of the number of associations per SNP in the raw dataset. Dashed line represents a threshold of 10 associations. **b.** A heatmap of Jaccard distances between pairs of phenotypes. Phenotypes are aligned according to the hierarchical clustering (HC). **c.** Same as (a) but for the clustered dataset. **d.** Comparison of numbers of associations per SNP before and after clustering of similar phenotypes. **e.** Violin plots representing minor allele frequency distribution for SNPs having either (1-5) or more than 5 associations in the clustered data.

**Figure 2.**
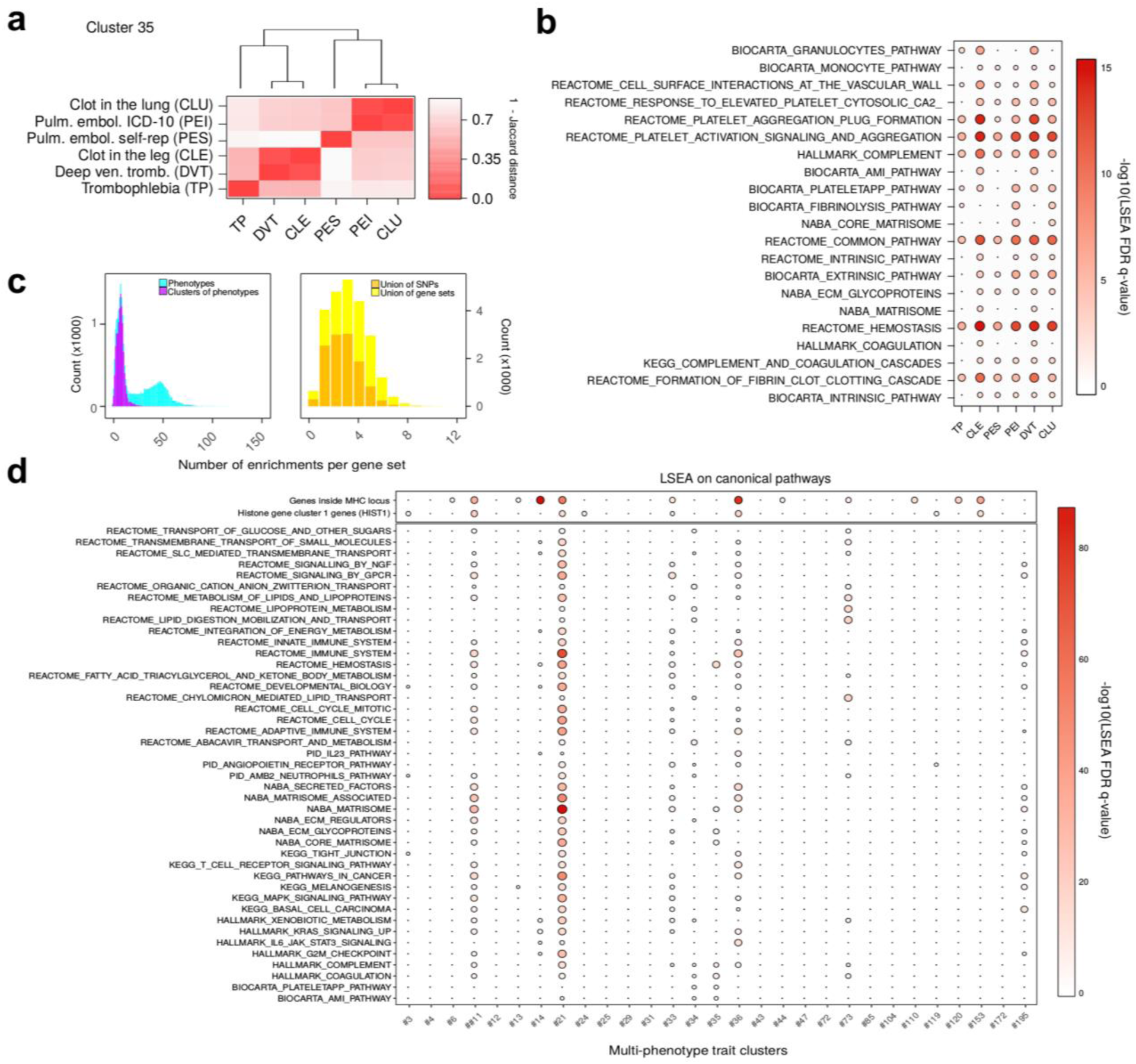
Multi-trait clusters share several key enriched molecular pathways. **(a).** A close-up view of the cluster 35 from Fig. 1c. Color indicates Jaccard distance in the space of significantly associated SNPs. **(b).** A circle-map representation of GSEA results for significantly associating SNPs for different phenotypes in cluster 35. Phenotypes are abbreviated as in (a). Circle size is proportional to the amount of overlapping loci; color intensity is proportional to the significance of enrichment (-log10(q-value)). **(c).** A histogram of numbers of enrichments per each gene set from the MSigDB C2 and H (canonical pathways and Hallmark) collections. Left panel, number of distinct associated phenotypes and phenotypes from distinct clusters; right panel, number of clusters showing significant enrichment based on SNPs shared by 2 or more phenotypes or based on gene sets shared by 2 or more phenotypes in each cluster. **(d).** A circle-map representation of GSEA results for common (shared by 2 or more phenotypes) associated SNPs for each multi-phenotype cluster. Only gene sets enriched in 2 or more clusters are shown (only top 100 enrichments for each cluster were chosen).

**Figure 3.**
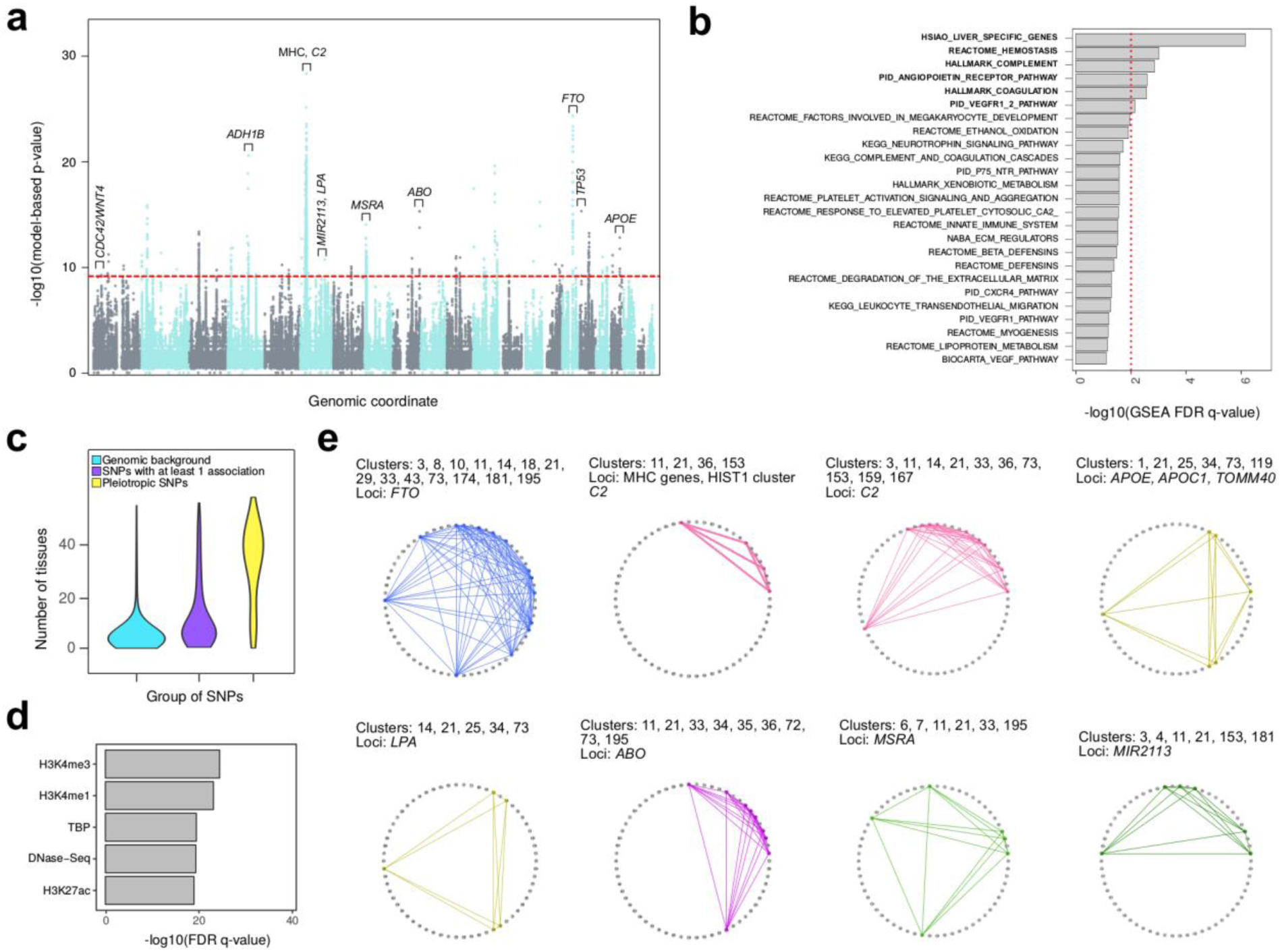
Loci with largest numbers of genetic associations belong to a limited set of most relevant molecular pathways. **a.** Manhattan plot of the pleiotropy p-value obtained from a Poisson model (see Methods). **b.** Results of the gene set enrichment analysis for a set of SNPs with genome-wide significant pleiotropy p-values. Gene sets with FDR Q-value < 0.01 are highlighted in bold. Only nominally significant hits are shown. **c.** Distributions of the number of tissues with significant eQTL signal for all SNPs, SNPs with at least 1 association, and pleiotropic SNPs (pleiotropy p-value < 5e-9). **d.** Results of the ChIP-Atlas enrichment analysis^22^ of the pleiotropic loci vs blood-derived genomic tracks (see Methods). Top-5 enriched tracks are shown. **e.** Network representation of associations for chosen groups of loci that are shared by a specific combination of clusters. Edge weight is proportional to the number of loci shared by a pair of nodes.

### Investigation of molecular pathways relevant to major trait clusters

We then focused on investigation of the genetic architecture of the major trait clusters. Many of the identified clusters comprised biologically similar complex traits or same phenotypes in different encoding (ICD-10 and self-reported disease status, etc.). As a way to decipher the molecular basis of traits and clusters, we used a modified version of the gene set enrichment analysis we call Locus Set Enrichment Analysis (LSEA) which accounts for linkage disequilibrium (LD) between SNPs and can work with few significantly associated SNPs for each trait (see Online methods for description of the method).

To focus solely on shared genetic architecture components in each cluster, we considered loci which were significantly associated with 2 or more phenotypes for each cluster. In contrast to other published methods^3,5^, LSEA approach sensitively identified enrichment for coagulation hallmark genes and specific coagulation pathways (e.g., intrinsic pathway, extrinsic pathway) for the cluster 35 comprising blood clotting and other related phenotypes (Figure 2a-b) accurately accounting for linkage between functionally related genes.

We next made functional annotation of all identified multi-phenotype clusters using the LSEA method. Cluster 21, comprising anthropometric traits, such as height, whole body weight, and properties of body parts and fat percentage, was enriched for diverse gene sets including extracellular matrix components and signalling pathways, such as GPCRs pathway, MAPK cascade and tyrosine kinases (see Extended Data Figure 4). For hypertension-related cluster 33, LSEA identified enrichments for platelet signalling, blood vessel formation, and hemostasis. Interestingly, we also observed highly significant enrichment for GPCR signaling pathway, which is concordant with investigation of GPCR signaling regulation in hypertension^17^. Yet another cluster comprised phenotypes characterized by high cholesterol level, with enriched gene sets predominantly related to lipid digestion, mobilization and transport, and small molecule metabolism. Genes from these gene sets were also overrepresented in the association signal of cluster 34, which included most heart pathologies, such as heart attack, angina, and acute myocardial infarction (see Extended Data Figure 7). Finally, cluster 36 related to autoimmune diseases such as asthma, hypo- and hyperthyroidism, and psoriasis, was expectedly enriched for immune system signaling pathways, e.g. cytokine-related ones (see Extended Data Figure 9).

Overall, we observed substantial overlap between gene sets enriched in major trait clusters. We identified 8393 gene sets that showed significant enrichment for at least 2 different phenotypes or clusters (760 of which belonged to the hallmark gene sets or canonical pathways set defined by Molecular Signatures Database (MSigDB)^18,19^. Of these, more than half had 50 or more phenotypes with enrichment signal (Figure 2c). We also investigated the enrichment statistics for another approach based on selection of gene sets enriched for at least 2 phenotypes in each cluster (Extended Data Figure 10). The results of such analysis were highly concordant with the ones obtained by using shared associated loci.

We then turned our attention to the analysis of gene sets which were highly enriched in the major trait clusters. To this end, we selected top-100 enriched gene sets for each cluster, and focused on gene sets having 2 or more enrichments after such filtering. Quite expectedly, we observed that most of the shared enriched gene sets belonged to the immune system, metabolism, and extracellular matrix (Figure 2d). Of these, hemostasis cascade elements, matrisome components, and xenobiotic metabolism genes were enriched in the largest number of trait clusters. We also assessed the enrichment of association signal inside two large regions of chromosome 6, the MHC region and the histone gene cluster 1, which is known to harbor multiple binding sites for the master regulator of MHC genes, *CIITA*^20^. We found 12 trait clusters to be significantly associated with variation inside the MHC locus, supporting its pivotal role in complex traits.

### Analysis of highly pleiotropic loci

As we observed a large amount of shared gene sets in our functional annotation studies (Figure 2d), we went on to systematically identify specific variants and loci involved in multiple complex traits and trait clusters. To this end, we first constructed a statistical model based on Poisson distribution that accounts for the randomly expected number of associated clusters for SNPs with different minor allele frequency (MAF), with such model accurately predicting the proportion of signal-harboring SNPs for each MAF bin. We used the constructed model to obtain the empirical *p-*value of multiple associations for each SNP given its MAF, and selected variants that showed genome-wide significance (P < 5 * 10^-9^); *i.e.* the ones involved in many more associations than expected by a random coincidence. Such filtering resulted in 2,000 genome-wide significant pleiotropic loci corresponding to 53 independent chromosomal regions. Pathway analysis of such pleiotropic loci identified strong enrichment for diverse immunity-related molecular pathways including complement system, chromosome maintenance, and hemostatic cascades.

We then went on to analyze the functional impact of the identified pleiotropic variants. We identified a very strong enrichment for cis-eQTLs (data collected from the Genotype Tissue Expression (GTEx) project^21^) in the set of pleiotropic variants (1953/2191 compared to 247,551/469,013, p-value = 1.2 * 10^-293^), suggesting that variants with high pleiotropic effect tend to have a significant influence on gene expression (Figure 3c). These results are in good concordance with the other reports on genetic pleiotropy in the UKB dataset^16^. We also assessed the prevalence of certain functional variant types in the previously defined set of pleiotropic variants. We detected significant overrepresentation of missense variants (46/2191 compared to 3073/469013, Fisher p-value = 2.4 * 10^-11^), as well as the other functionally important variant classes, among the pleiotropic variants compared to variants associated with at least one trait (see Extended Data Figure 11). Furthermore, we identified a strong enrichment for open chromatin and promoter marks (H3K4Me3, H3K27Ac, TBP, DNAse hypersensitivity) in the set of pleiotropic loci using genomic track enrichment analysis (ChIP-Atlas^22^, Figure 3d).

We then set off to characterize the specific sets of loci shared by different clusters. Firstly, we investigated the overall structure of the pleiotropy network represented as weighted graph with nodes corresponding to trait clusters and edges corresponding to shared pleiotropic loci defined above. 62 trait clusters shared at least one pleiotropic locus with the others (see Extended Data Figure 12), with each pair of clusters shared 2.4 loci on average (see Extended Data Figure 13). The anthropometric trait-related cluster (21) shared at least one pleiotropic locus with all the others, while several other clusters (11, 33, and 36) shared loci with a large number of clusters.

We then continued by selecting highly connected subgraphs of the pleiotropy network (combinations of nodes sharing a specific set of loci) (see Methods). While the most noticeable sets of shared pleiotropic loci belonged to the MHC complex, we found numerous interesting examples of shared groups of loci (shown in Figure 3e). As expected, larger combinations of trait clusters tended to share fewer common loci (see Extended Data Figure 14).

One of the most notable loci included the *FTO* gene, which was found to be associated with 14 clusters. This finding is concordant with previous reports suggesting this gene to be connected to obesity and many obesity-related comorbidities^23^. Variants in the *FTO* gene affecting body composition and metabolic rates, are likely to increase risk of diabetes (clusters 14, 34), vascular problems (33), and lipid metabolism disorders (73). Another interesting single-gene pleiotropic locus comprised the *MSRA* gene associated with trait clusters related to anxiety (6), neuroticism (11), and sleep duration (7). This gene has already been linked to neuroticism^24^. We also found the *LPA* gene to be connected to clusters comprising diverse heart problems and increased cholesterol level, including myocardial infarction and chronic ischemic heart disease (belonging to cluster 34), high cholesterol level (73), and ezetimibe intake (24). The relationship between SNPs in lipoprotein(a) [Lp(a)] and major adverse cardiovascular (MACE) events has also previously been reported^25^.

Among the other genes, *ABO* was found to be associated with trait clusters related to a wide variety of vascular phenotypes such as warfarin treatment (72), blood pressure dysregulation (33), embolism (35), and heel bone mineral density (195). While the role of *ABO* in thrombosis has been excessively studied^26^, the overall significance of this locus to the majority of cardiovascular phenotypes has not been previously investigated. Yet another group of loci included *APOE* and *TOMM40* genes associated with body mass index (21), diverse lipid-metabolism associated phenotypes (73), Alzheimer’s disease (25). Observed data is quite convergent with studies of *APOE* and *TOMM40* alleles causing Alzheimer’s disease^27^, and further support a solid link between lipid metabolism and diverse complex phenotypes.

One of the most interesting pleiotropic loci included the *MIR2113* gene encoding a microRNA of unknown function. This locus is associated with several phenotypes related to cognitive characteristics, including college or university degree qualifications (cluster 11) and fluid intelligence score (cluster 153), as well as ion concentrations (sodium in urine (181), potassium in urine (153)). Previously, *MIR2113* was shown to play a role in memory processes^28^. However, the exact mechanism of its function has never been reported. Our data on its involvement in maintenance of ionic balance, together with a strong enrichment for potassium channel genes among its predicted targets according to miRBase (Extended Data Figure 15), suggest that *MIR2113* regulates cognitive processes through regulation of ion channel gene expression, which is important for understanding the development of cognition-related traits.

## Discussion

Large-scale genetic datasets, such as the UK Biobank or the Genome Aggregation Database (gnomAD)^29^ allow the first ever look into the ‘Holy Grail’ of human genetics research, *i.e.* the map of genotype-phenotype relationships. In our study, we aimed at identification of highly pleiotropic loci in the human genome and dissecting the key genetic architecture components relevant to human complex traits. The UKB genetic dataset contains a massive amount of similar and/or related traits which are commonly separated into several major domains^16,30^. To overcome this limitation and group traits that share a significant proportion of their genetic architecture independently of their arbitrary classification, we used a hierarchical clustering approach using the simple Jaccard distance metric calculated using sets of significantly associated variants. While such an approach could misclassify phenotypes with low signal-to-noise ratio, it efficiently clusters similar and identical phenotypes. The results of such clustering approach are concordant with the other efforts to computationally classify traits in the UKB dataset^12,16^. Notably, our clustering procedure identified close relationships between traits that have been demonstrated as comorbid to each other. For instance, deep venous thrombosis and pulmonary embolism combine in one cluster (35) with diverse thromboembolisms. It was clearly shown that deep venous thrombosis significantly increases the risk of pulmonary embolism^25^. Grouping of asthma, diabetes, and increased risk of nasal polyps (cluster 14) is also consistent with known data^31,32^. Therefore, our results are in good concordance with existing biological and clinical knowledge.

One of the major results presented in this work is the comprehensive functional annotation of GWAS signal for individual traits and trait clusters. To perform such functional annotation using a limited set of significant variants, we’ve constructed an LD-aware algorithm for GSEA (LSEA), which allowed us to identify crucial molecular pathways involved in multiple groups of phenotypes. For example, we discovered multiple associations for genes involved in innate and adaptive immunity (Figure 2d), as well as an overwhelming enrichment for variants inside the HLA locus. These results provide additional evidence that variation in genes and proteins involved in immune system functioning affects a large number of developmental and physiological processes. Similar observations have been made previously using site-level PheWAS^33^. Interestingly, we also discover high degree of pleiotropy for the loci inside the *HIST1* gene cluster, which is known to harbor multiple binding sites for the *CIITA* transactivator protein that regulates MHC gene expression^34^. We also observe stronger association with traits from the smoking-related cluster (24) for variants inside the *HIST1* gene cluster than for the MHC genes themselves. This association signal provides new mechanistic insights into the previously reported association of the histone genes with increased risk of squamous cell lung carcinoma^35^. Another set of genes involved in multiple complex trait clusters comprises the matrisome components. Matrisome genes are involved in the highest amount of individual phenotypes and clusters, including the anthropometric trait cluster 21. This is not surprising, regarding functional significance of ECM proteins in governing body compositions and their previously reported role in diverse diseases^36^.

Deciphering the mechanisms of complex traits and their co-morbidity is a capstone for GWAS data interpretation and further clinical application of the results^37^. In this regard, Mendelian randomization (MR) approaches could be quite useful when analyzing chains of causal effects between genetic variants and their epistatic interactions that lead to phenotypic manifestation. MR approaches have successfully been applied to investigation of genetic pleiotropy and identification of variants influencing multiple traits^38^. In our study, we applied a different statistical approach to investigate multiple association in genome-wide data. Noteworthy, the results of our network analysis of pleiotropic interactions (Figure 3) corroborate and expand previous MR-based findings. For example, we report much higher multiplicity of genetic associations for the *FTO* locus, comprising breast cancer, heel bone mineral density, anthropometric traits and sleep duration. We report additional associations for the diabetes-related *GCKR* gene, such as gout, high cholesterol levels and narcolepsy. We also expand the list of associated phenotypes for the *APOE* gene, adding vascular/heart problems (chronic ischemic heart disease and angina) and pulse rate. Among the sets of pleiotropic loci we found the *SLC39A8* gene associated with osteoporosis, as also reported previously^39^. The same locus is reported to be associated with another osteoporosis- and osteoarthritis-related traits that were also investigated in a recent meta-analysis of UKB data^40^. These include wait-hip ratio, BMI, body fat percentage, strenuous sport, alcohol intake, and prospective memory result. The exact relationships between these traits, though, are still to be investigated.

We identify several key properties of the pleiotropic variation. First, SNPs with high amount of associated traits and clusters tend to be common (Figure 1e, Extended Data Figure 3). This is unexpected given that variants with larger effect sizes tend to be rare due to negative selection^8^. However, low degree of pleiotropy for rare variants might be explained by sample size limitations. Secondly, pleiotropic loci identified in this study are significantly enriched in open chromatin regions bearing H3K4Me3 and H3K27Ac histone modifications as identified by ChIP-Atlas search^22^ (Figure 3d). This is consistent with previous observations made for immune system^41^. Third, variants with high degree of pleiotropy have higher functional impact: these are enriched for both protein-altering missense variants and broader range eQTLs (Figure 3c). All of the aforementioned properties of pleiotropic loci are in very good concordance with the results of other global overviews of pleiotropy made using UKB data^15,16^.

Overall, our approach allowed us to detect multiple novel interrelations between complex traits, which can shed light on common molecular patterns driving the human phenome. Further investigations would help to determine causal relationships between key genes and molecular pathways that give rise to multigenic disease. Investigation of all possible genetic interactions between common and specific risk factors should help to decipher sophisticated molecular mechanisms behind complex traits.

## Acknowledgements

We thank Mykyta N. Artomov for useful comments and suggestions. The authors declare no conflict of interest.

## Methods

### Data acquisition and filtering criteria

We acquired data from the Benjamin Neale’s lab website (UK Biobank GWAS results imputed v2, downloaded at 2017-10-04; http://www.nealelab.is/uk-biobank). We focused our attention only on phenotypes with significant non-zero partitioned heritability estimates (p < 0.05, estimates provided by the authors of the dataset). After obtaining the list of heritable phenotypes we retained genome-wide significant variants for each phenotype (p-value < 5 * 10^-9^). Association summary statistics for each variant for each trait were then merged into a singly matrix for further analysis.

### Clustering of traits

To group similar traits in space of pre-selected associated variants, we utilized the Jaccard distance metric calculated for each pair of traits as follows:

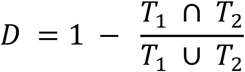

where *D* is the distance between two phenotypes, and *T*_*1*_ and *T*_*2*_ are sets of associated SNPs for two traits. We then performed hierarchical clustering together with the analysis of silhouettes to determine the most appropriate number of clusters. We then calculated the number of associated trait clusters for each variant. To this end, we considered an SNP to be associated with a cluster if this SNP is associated with at least one trait inside this cluster.

### Description of the LSEA method

We implemented a novel Locus Set Enrichment Analysis (LSEA) method for gene set enrichment analysis of GWAS data. LSEA operates on the sets of pre-selected significant SNPs or summary statistics file for each trait. At first stage, LSEA groups associated SNPs into individual genomic loci. First, variants are clustered by LD using the clumping method in PLINK^42^. The resulting groups of SNPs are transformed to an interval list based on the leftmost and rightmost coordinate of variants in each clump. The resulting intervals are then merged if these overlap by more than 70%, or span different parts of the same gene, and intervals overlapping the MHC region or the *HIST1* gene cluster (defined as chr6:28,866,528-33,775,446 and chr6:25,000,528-28,000,446, respectively, based on hg19 human genome annotation) are omitted, as suggested previously^3^. The resulting intervals are then intersected with the pre-computed universe of all possible loci bearing GWAS signal.

To construct such universe, we merged all loci that were identified for the whole UKB dataset. We then transformed all of the curated gene sets obtained from the MsigDB database (http://software.broadinstitute.org/gsea/msigdb/annotate.jsp) the following way: for each interval *i* in the intervals universe, we assigned *i* to a gene set if at least one gene in the gene set overlaps *i*. Such method efficiently corrects for functionally related genes that are genetically linked, as intervals spanning several genes belonging to each gene set are counted only once.

The enrichment statistic is computed for each gene set as follows:

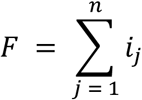

where *n* is the number of intervals in the query set, *i*_*j*_ = 1 if interval *j* belongs to a gene set of interest, and *i*_*j*_ = 0 otherwise. Enrichment p-value is then computed from hypergeometric distribution. The resulting p-values are then adjusted for multiple comparisons using the Holm FDR correction.

### Gene set enrichment analysis of individual traits and clusters

We used the LSEA method to analyze gene set enrichment for all individual phenotypes and trait clusters. We used two complementary approaches to make functional annotations of multi-trait clusters. The first approach was based on using variants that are significantly associated with at least 2 of the traits in each cluster (termed “union of SNPs” on Figure 2c). For the second approach, we used LSEA results for all individual phenotypes in each cluster, and retained only gene sets that were significantly enriched for 2 or more traits in a cluster (termed “union of gene sets” on Figure 2c.

As the HLA locus and the *HIST1* gene cluster were omitted during gene set analysis with LSEA, we analyzed the overrepresentation of variants inside these loci separately. To this end, we created sets of intervals for the *HIST1* and HLA regions in a way analogous to the LSEA method. We then obtained the overrepresentation p-values using hypergeometric distribution, with the test statistic being equal to the number of intervals in the query that overlap the regions of interest.

### Identification of pleiotropic genetic variants

To assess the phenome-wide significance and identify key pleiotropic variants, we constructed a model to account for the null expectation of random overlap between association signals. If the association signals are randomly distributed across the genome for two individual phenotypes (or clusters), the number of per-SNP associations should follow a Poisson distribution in a manner similar to the sequencing read coverage^43^. As we noted that SNPs with different MAF differ in the average number of associations (Figure 1e, Extended Data Figure 3), we divided the variants into 500 bins based on MAF ranged from 0 to 0.5. We then calculated the mean number of associations for variants in each bin (λ, see Extended Data Figure 16). This parameter was then used to calculate the phenome-wide significance p-value:

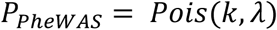

where *k* is the number of traits or trait clusters associated with a given variant. Variants with p-value < 5 * 10^-9^ were selected as pleiotropic SNPs and used for further analyses.

### Overrepresentation analysis for functional variant classes

To analyze the overrepresentation of cis-eQTLs, we used the GTEx data (REF). Sets of variants of interest (i.e., pleiotropic variants and variants with at least 1 association) were overlapped with significant cis-eQTLs (p < 5 * 10^-8^). Overrepresentation of cis-eQTLs in the set of pleiotropic SNPs was assessed by comparison of the resulting proportion of significant eQTLs using the hypergeometric distribution.

We also annotated all SNPs from the UKB dataset with SnpEff^44^ and separated the variants by their functional type (e.g., missense, intronic, etc. (the distributions are shown in Extended Data Figure 14)). We then compared the proportions of certain functional classes of variants in the sets of pleiotropic SNPs and SNPs bearing at least one association. The proportions were compared using the hypergeometric test.

To analyze the enrichment of epigenetic marks and chromatin states in the set of pleiotropic loci, we used the previously described ChIP-Atlas server^22^. For this analysis, sets of pleiotropic variants were converted to an interval list, and the resulting intervals were used as the input for ChIP-Atlas. To narrow down the search space, we narrowed the analysis down to a set of blood-derived genomic tracks. For each type of epigenetic mark, we retained the minimal enrichment p-value for further analysis.

### Network analysis of pleiotropic loci

To group loci that share sets of associated clusters, we constructed a network of genetic associations for the identified pleiotropic loci. To this end, we created a graph representation of the data, with nodes corresponding to individual clusters and edge weight between a pair of adjacent nodes corresponding to the number of shared independent loci. To search for individual fully connected subgraphs of this network (*i.e.*, nodes sharing the same sets of loci) we used a simple deterministic algorithm. At the first stage, we identified all possible large combinations of nodes connected by an individual pleiotropic locus by iterating over edges of the graph. We then merged loci that share the same combinations of associated nodes.

### Data and source code availability

All data and scripts pertinent to the analysis presented here, as well as the source code for LSEA, are available through GitHub (https://github.com/bioinf/ukb_phewas).

## References

1. Mills, M. C. & Rahal, C. A scientometric review of genome-wide association studies. Commun. Biol. 2, (2019).

2. Mahajan, A. et al. Fine-mapping type 2 diabetes loci to single-variant resolution using high-density imputation and islet-specific epigenome maps. Nat. Genet. 50, 1505–1513 (2018).

3. Genetic Investigation of ANthropometric Traits (GIANT) Consortium et al. Biological interpretation of genome-wide association studies using predicted gene functions. Nat. Commun. 6, (2015).

4. de Leeuw, C. A., Mooij, J. M., Heskes, T. & Posthuma, D. MAGMA: Generalized Gene-Set Analysis of GWAS Data. PLOS Comput. Biol. 11, e1004219 (2015).

5. Watanabe, K., Taskesen, E., van Bochoven, A. & Posthuma, D. Functional mapping and annotation of genetic associations with FUMA. Nat. Commun. 8, (2017).

6. GTEx Consortium et al. Using an atlas of gene regulation across 44 human tissues to inform complex disease- and trait-associated variation. Nat. Genet. 50, 956–967 (2018).

7. Sudlow, C. et al. UK Biobank: An Open Access Resource for Identifying the Causes of a Wide Range of Complex Diseases of Middle and Old Age. PLOS Med. 12, e1001779 (2015).

8. Zeng, J. et al. Signatures of negative selection in the genetic architecture of human complex traits. Nat. Genet. 50, 746–753 (2018).

9. Khera, A. V. et al. Genome-wide polygenic scores for common diseases identify individuals with risk equivalent to monogenic mutations. Nat. Genet. 50, 1219–1224 (2018).

10. Canela-Xandri, O., Rawlik, K. & Tenesa, A. An atlas of genetic associations in UK Biobank. Nat. Genet. 50, 1593–1599 (2018).

11. McInnes, G. et al. Global Biobank Engine: enabling genotype-phenotype browsing for biobank summary statistics. Bioinformatics (2018). doi:10.1093/bioinformatics/bty999

12. Gaspar, H. A., Hübel, C., Coleman, J. R., Hanscombe, K. B. & Breen, G. Navigome: Navigating the Human Phenome. bioRxiv (2018). doi:10.1101/449207

13. Schmidt, A. F. et al. Phenome-wide association analysis of LDL-cholesterol lowering genetic variants in PCSK9. bioRxiv (2018). doi:10.1101/329052

14. Verbanck, M., Chen, C.-Y., Neale, B. & Do, R. Detection of widespread horizontal pleiotropy in causal relationships inferred from Mendelian randomization between complex traits and diseases. Nat. Genet. 50, 693–698 (2018).

15. Jordan, D. M., Verbanck, M. & Do, R. The landscape of pervasive horizontal pleiotropy in human genetic variation is driven by extreme polygenicity of human traits and diseases. bioRxiv (2019). doi:10.1101/311332

16. Watanabe, K. et al. A global view of pleiotropy and genetic architecture in complex traits. bioRxiv (2018). doi:10.1101/500090

17. Brinks, H. L. & Eckhart, A. D. Regulation of GPCR signaling in Hypertension. Biochim. Biophys. Acta BBA - Mol. Basis Dis. 1802, 1268–1275 (2010).

18. Liberzon, A. et al. Molecular signatures database (MSigDB) 3.0. Bioinformatics 27, 1739–1740 (2011).

19. Liberzon, A. et al. The Molecular Signatures Database Hallmark Gene Set Collection. Cell Syst. 1, 417–425 (2015).

20. Wong, D. et al. Genomic mapping of the MHC transactivator CIITA using an integrated ChIP-seq and genetical genomics approach. Genome Biol. 15, (2014).

21. Lonsdale, J. et al. The Genotype-Tissue Expression (GTEx) project. Nat. Genet. 45, 580–585 (2013).

22. Oki, S. et al. ChIP-Atlas: a data-mining suite powered by full integration of public ChIP-seq data. EMBO Rep. 19, e46255 (2018).

23. Loos, R. J. F. & Yeo, G. S. H. The bigger picture of FTO—the first GWAS-identified obesity gene. Nat. Rev. Endocrinol. 10, 51–61 (2014).

24. Boutwell, B. et al. Replication and characterization of CADM2 and MSRA genes on human behavior. Heliyon 3, e00349 (2017).

25. Lee, J. S. et al. Deep Vein Thrombosis in Patients with Pulmonary Embolism: Prevalance, Clinical Significance and Outcome. Vasc. Spec. Int. 32, 166–174 (2016).

26. Long, Z., Du, Y., Li, H. & Han, B. Polymorphism of the *ABO* gene associate with thrombosis risk in patients with paroxysmal nocturnal hemoglobinuria. Oncotarget 8, (2017).

27. Nishimura, A. et al. Characterization of APOE and TOMM40 allele frequencies in the Japanese population. Alzheimers Dement. Transl. Res. Clin. Interv. 3, 524–530 (2017).

28. Andrews, S. J., Das, D., Anstey, K. J. & Easteal, S. Association of *AKAP6* and *MIR2113* with cognitive performance in a population-based sample of older adults: *AKAP6* and *MIR2113* in cognitive decline. Genes Brain Behav. 16, 472–478 (2017).

29. Karczewski, K. J. et al. Variation across 141,456 human exomes and genomes reveals the spectrum of loss-of-function intolerance across human protein-coding genes: Supplementary Information. bioRxiv (2019). doi:10.1101/531210

30. Ge, T., Chen, C.-Y., Neale, B. M., Sabuncu, M. R. & Smoller, J. W. Phenome-wide heritability analysis of the UK Biobank. PLOS Genet. 13, e1006711 (2017).

31. Forno, E. Asthma and diabetes: Does treatment with metformin improve asthma?: Editorial. Respirology 21, 1144–1145 (2016).

32. Zhang, Z. et al. The effect of diabetes mellitus on chronic rhinosinusitis and sinus surgery outcome: Impact of diabetes on chronic rhinosinusitis. Int. Forum Allergy Rhinol. 4, 315–320 (2014).

33. Verma, A. et al. Phenome-Wide Association Study to Explore Relationships between Immune System Related Genetic Loci and Complex Traits and Diseases. PLOS ONE 11, e0160573 (2016).

34. Wong, D. et al. Genomic mapping of the MHC transactivator CIITA using an integrated ChIP-seq and genetical genomics approach. 15 (2014).

35. Rosenberger, A. et al. Gene-set meta-analysis of lung cancer identifies pathway related to systemic lupus erythematosus. PLOS ONE 12, e0173339 (2017).

36. Hynes, R. O. & Naba, A. Overview of the Matrisome--An Inventory of Extracellular Matrix Constituents and Functions. Cold Spring Harb. Perspect. Biol. 4, a004903–a004903 (2012).

37. Gratten, J. & Visscher, P. M. Genetic pleiotropy in complex traits and diseases: implications for genomic medicine. Genome Med. 8, (2016).

38. Pickrell, J. K. et al. Detection and interpretation of shared genetic influences on 42 human traits. Nat. Genet. 48, 709–717 (2016).

39. Song, J. et al. MicroRNA-488 regulates zinc transporter SLC39A8/ZIP8 during pathogenesis of osteoarthritis. J. Biomed. Sci. 20, 31 (2013).

40. Zengini, E. et al. Genome-wide analyses using UK Biobank data provide insights into the genetic architecture of osteoarthritis. Nat. Genet. 50, 549–558 (2018).

41. Farh, K. K.-H. et al. Genetic and epigenetic fine mapping of causal autoimmune disease variants. Nature 518, 337–343 (2015).

42. Purcell, S. et al. PLINK: A Tool Set for Whole-Genome Association and Population-Based Linkage Analyses. Am. J. Hum. Genet. 81, 559–575 (2007).

43. Evans, S. N., Hower, V. & Pachter, L. Coverage statistics for sequence census methods. BMC Bioinformatics 11, 430 (2010).

44. Cingolani, P. et al. A program for annotating and predicting the effects of single nucleotide polymorphisms, SnpEff: SNPs in the genome of Drosophila melanogaster strain w ^1118^; iso-2; iso-3. Fly (Austin) 6, 80–92 (2012).

