## Extended data figures 1-16 for "Phenome-wide search for pleiotropic loci highlights key genes and molecular pathways for human complex traits"

**Article type:** Analysis

**Keywords:** phenome-wide association study, pleiotropy, molecular pathway, complex trait

### Extended Data

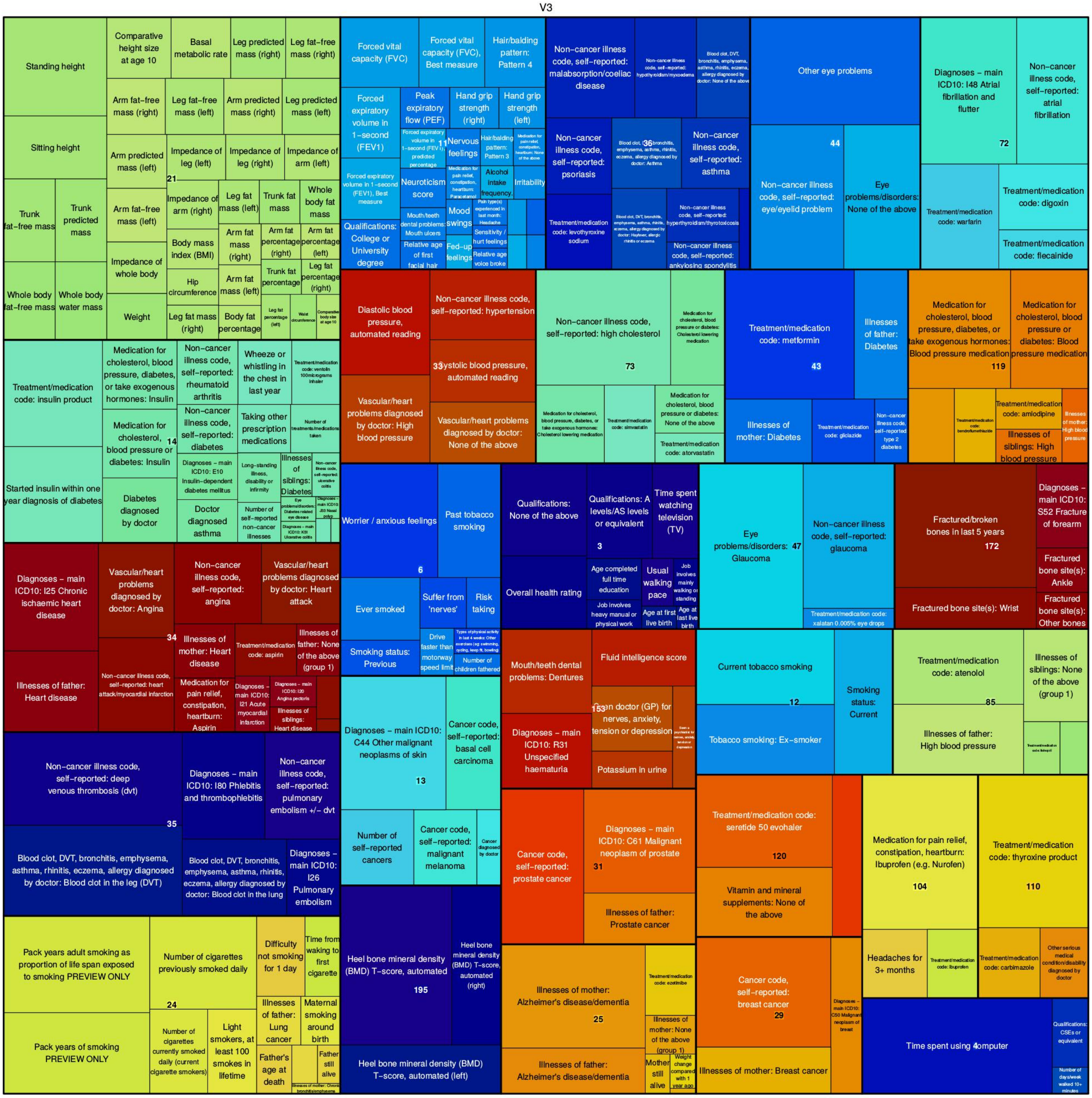


**Extended Data Figure 1.** Voronoi treemaps of multi-phenotype clusters defined by hierarchical clustering of the significant SNPs. Colour blocks correspond to individual cluasters. Numbers are equal to the number of cluster used throughout the work.

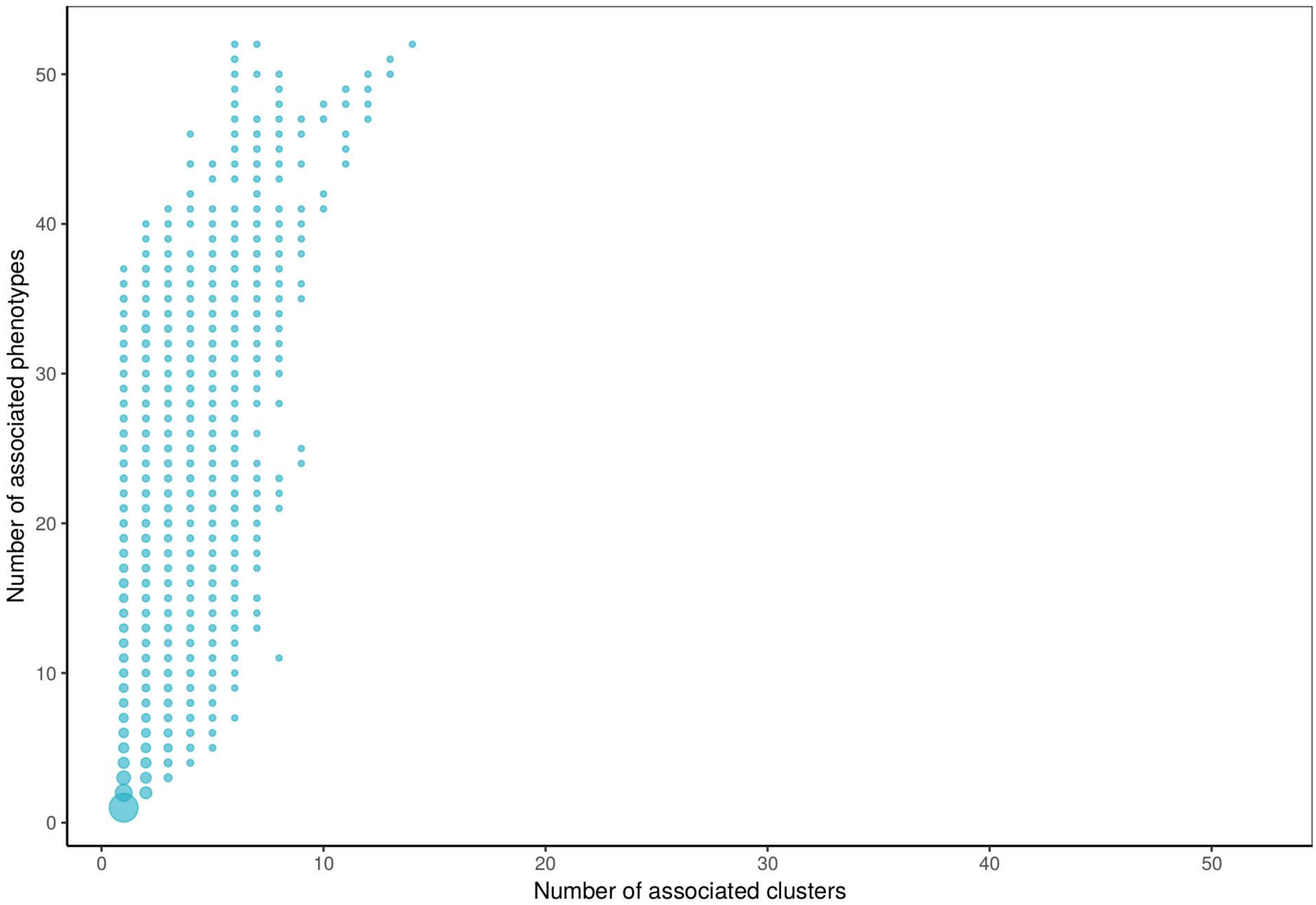


**Extended Data Figure 2.** A scatterplot of the per-SNP number of associated phenotypes and phenotype clusters. Dot size is proportional to the log10 (number of SNPs).

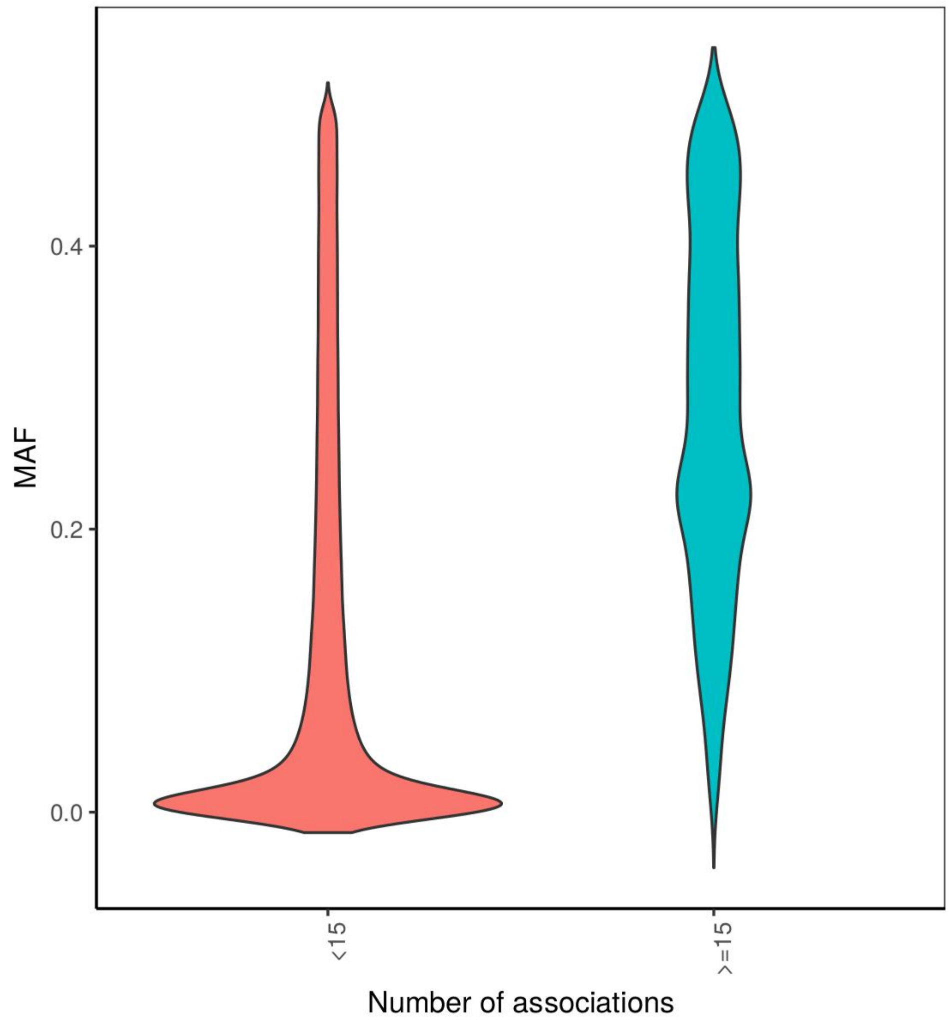


**Extended Data Figure 3.** Violin plots representing minor allele frequency distribution for SNPs having either (1-14) or more than 14 associations in the non-clustered data.

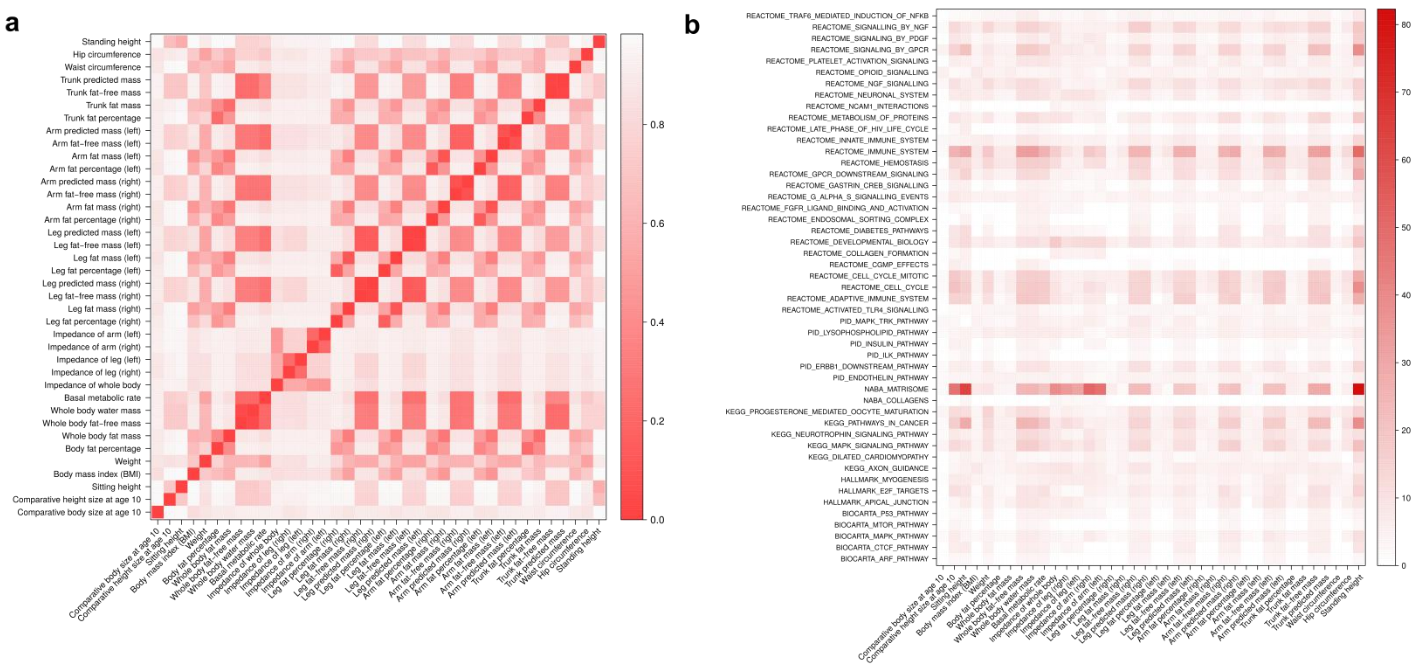


**Extended Data Figure 4.** Overview of the phenotype cluster #21 (anthropometric data). **a.** A fragment of the distance matrix. **b.** top 100 gene sets enriched for at least 2 phenotypes. Order of the phenotypes on the right is similar to the one on the left.

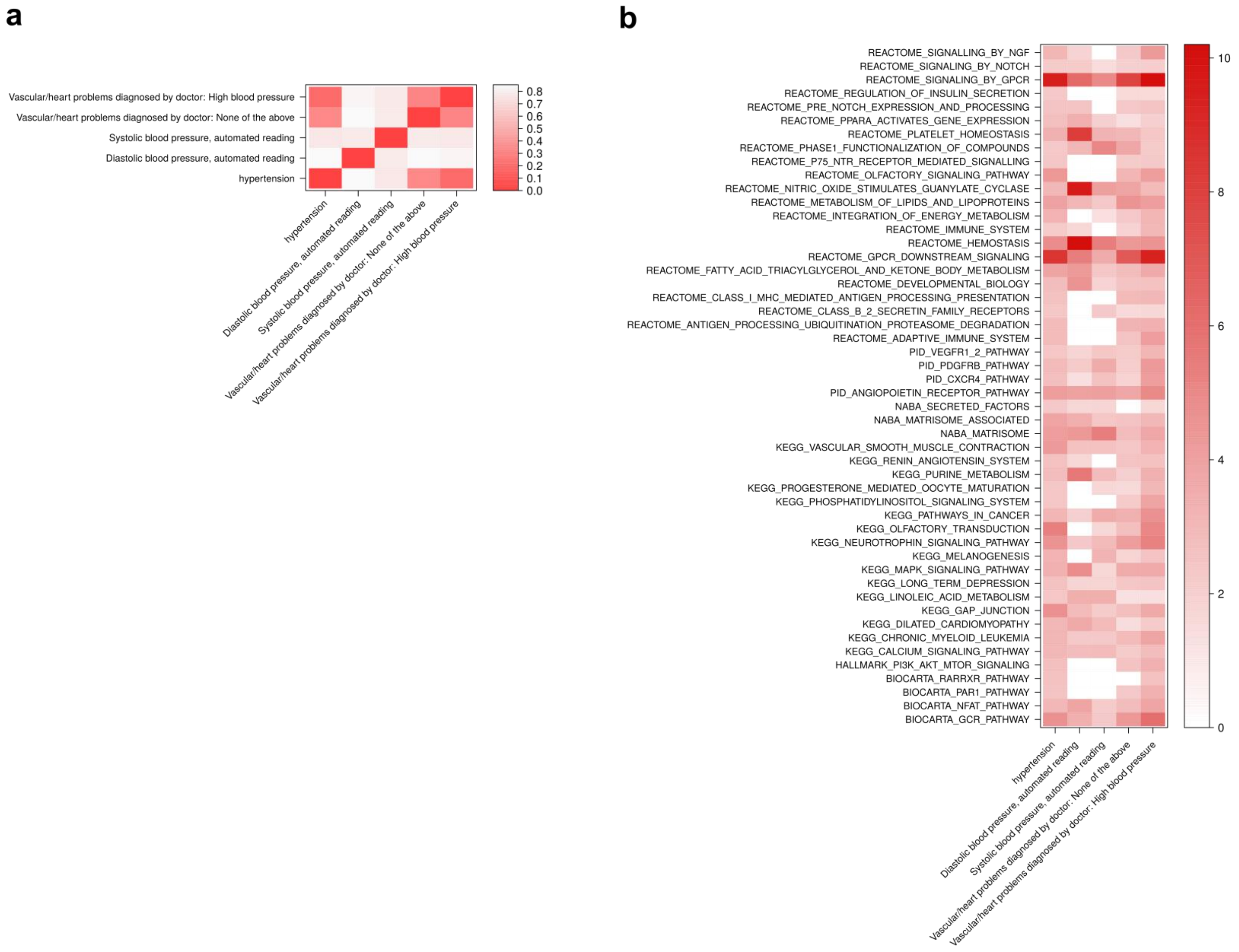


**Extended Data Figure 5.** Overview of the phenotype cluster #21 (anthropometric data). Left, a fragment of the distance matrix. Right, top 100 gene sets enriched for at least 2 phenotypes. Order of the phenotypes on the right is similar to the one on the left.

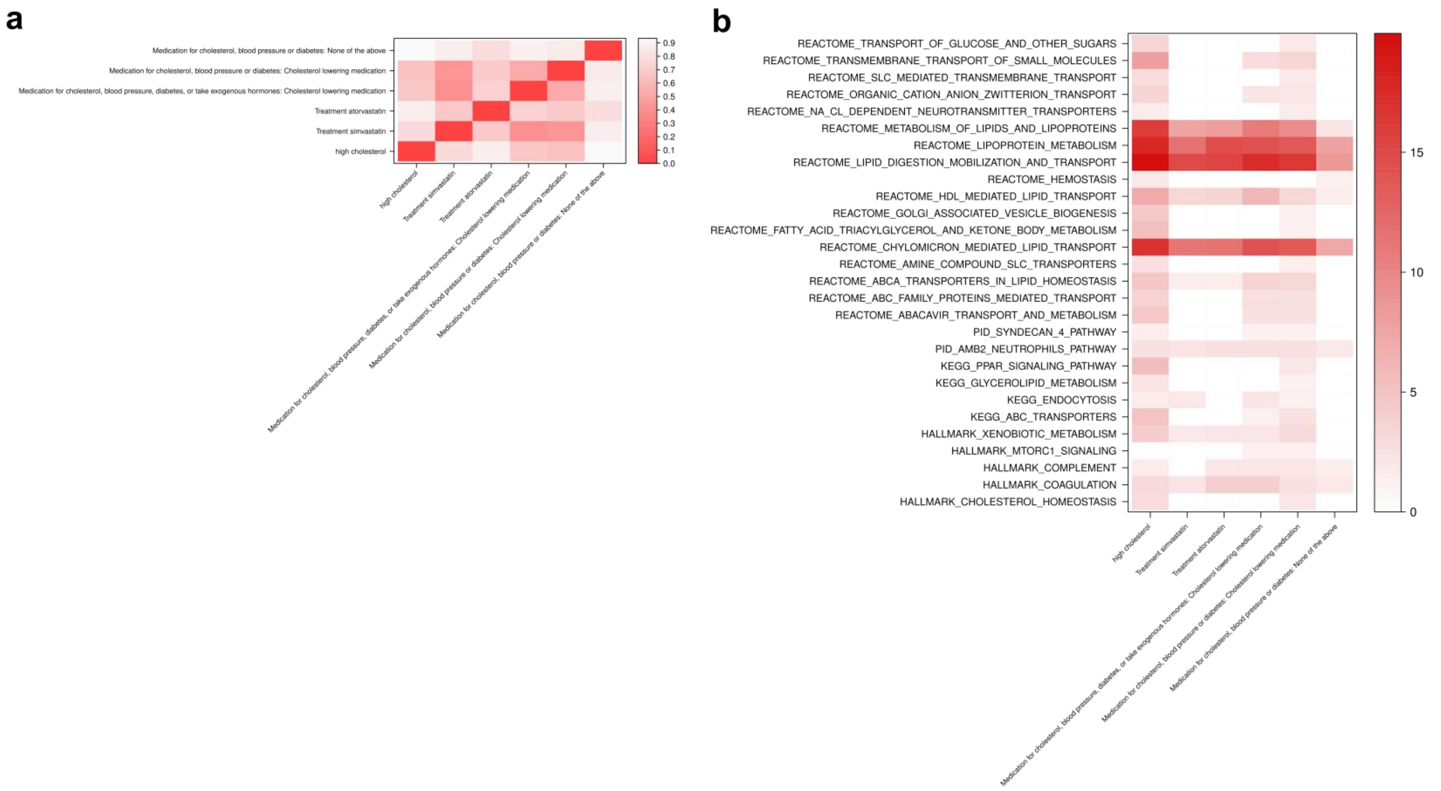


**Extended Data Figure 6.** Overview of the phenotype cluster #21 (anthropometric data). Left, a fragment of the distance matrix. Right, top 100 gene sets enriched for at least 2 phenotypes.

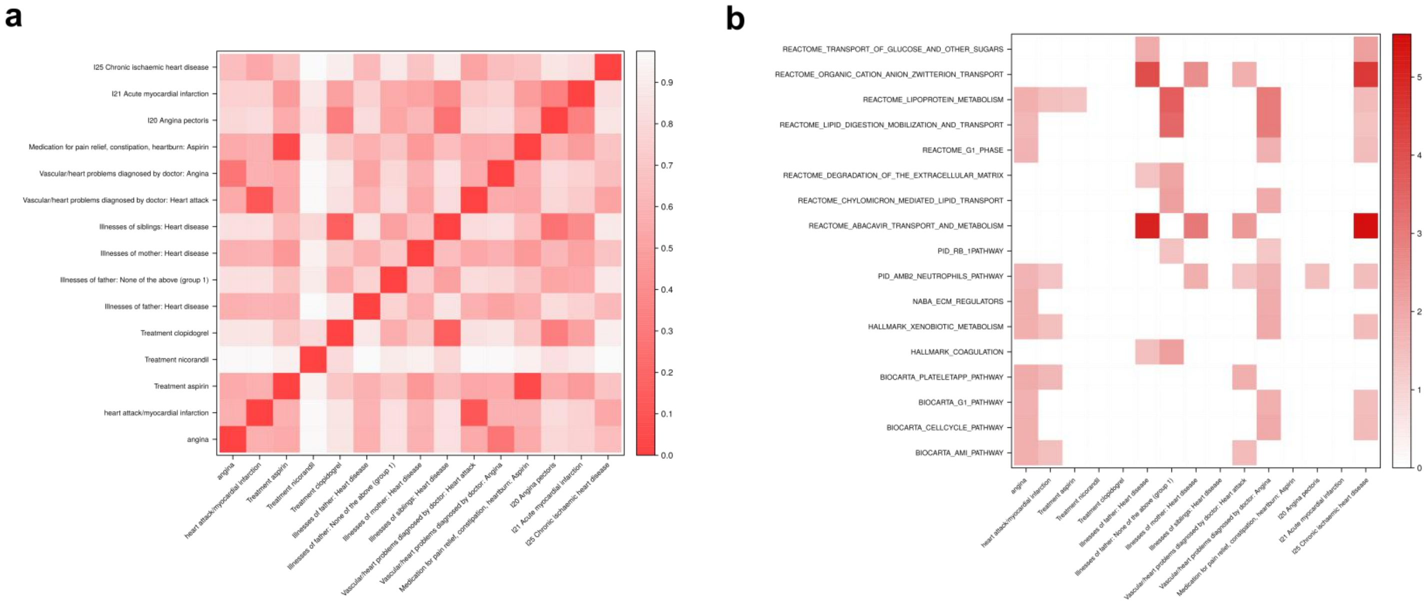


**Extended Data Figure 7.** Overview of the phenotype cluster #21 (anthropometric data). Left, a fragment of the distance matrix. Right, top 100 gene sets enriched for at least 2 phenotypes.

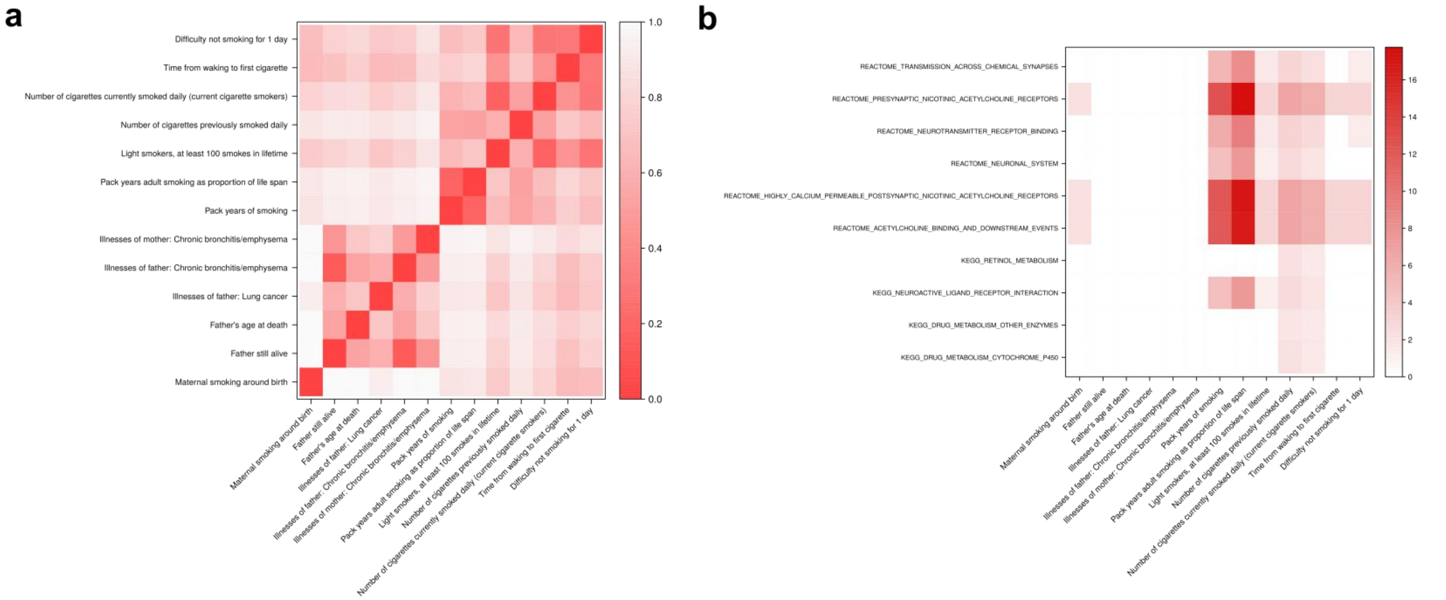


**Extended Data Figure 8.** Overview of the phenotype cluster #21 (anthropometric data). Left, a fragment of the distance matrix. Right, top 100 gene sets enriched for at least 2 phenotypes.

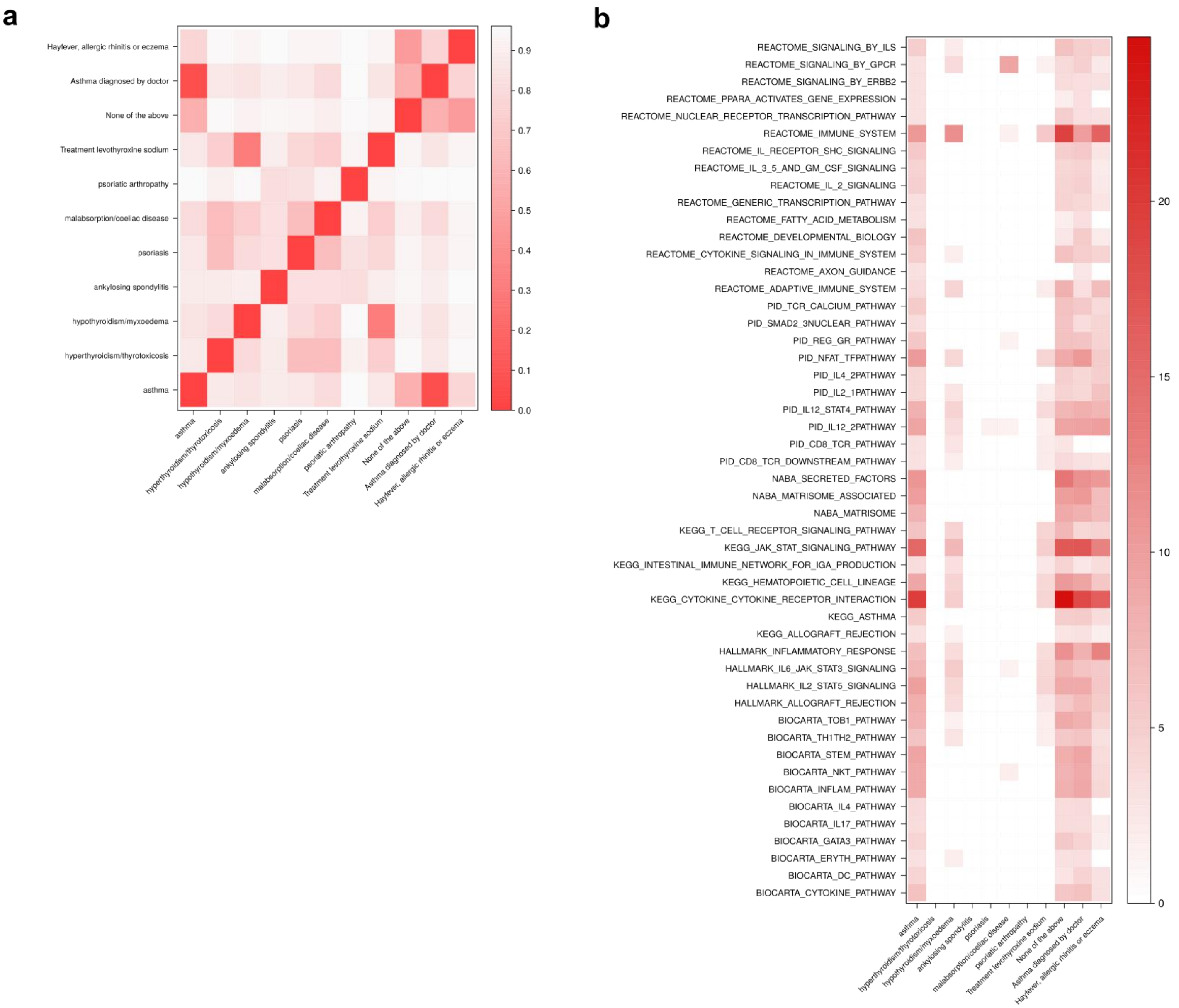


**Extended Data Figure 9.** Overview of the phenotype cluster #21 (anthropometric data). Left, a fragment of the distance matrix. Right, top 100 gene sets enriched for at least 2 phenotypes.

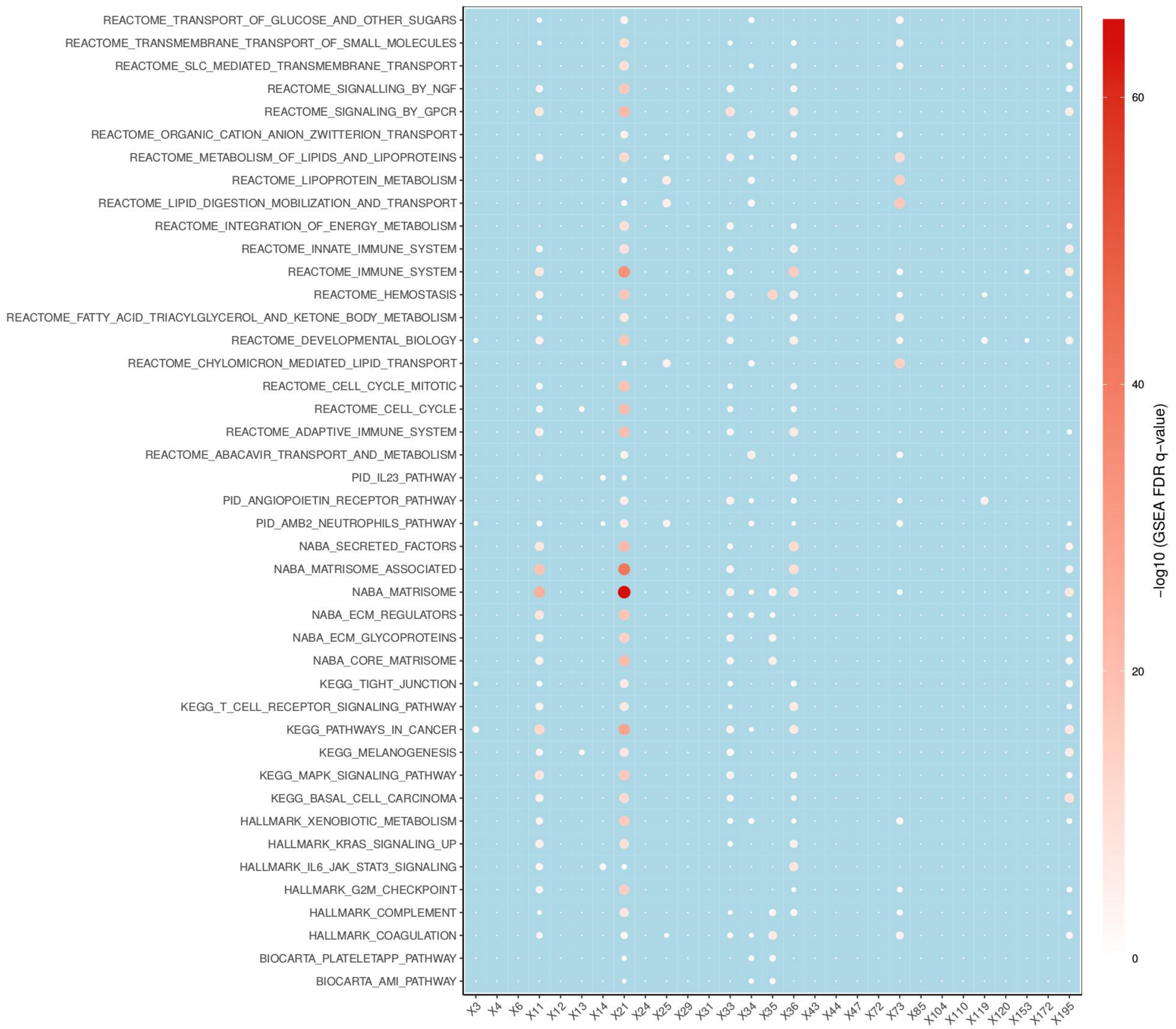


**Extended Data Figure 10.** A graphical representation of gene set enrichment analysis results for each trait cluster. Shown are gene sets that are enriched for at least 2 phenotypes in a cluster. P-values correspond to the second lowest p-value for all phenotypes in each cluster.

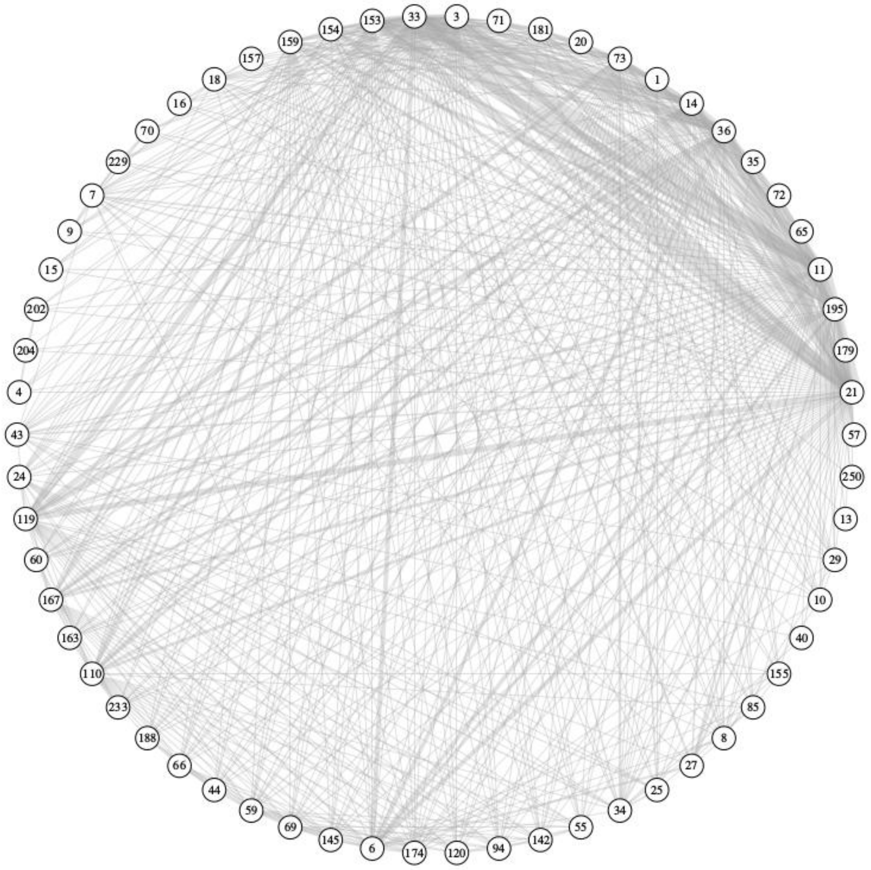


**Extended Data Figure 11.** A complete network of pleiotropic interactions between phenotype clusters. Each edge coreesponds to one or more shared pleiotropic loci. Edge weight represents the number of shared loci for a pair of clusters.

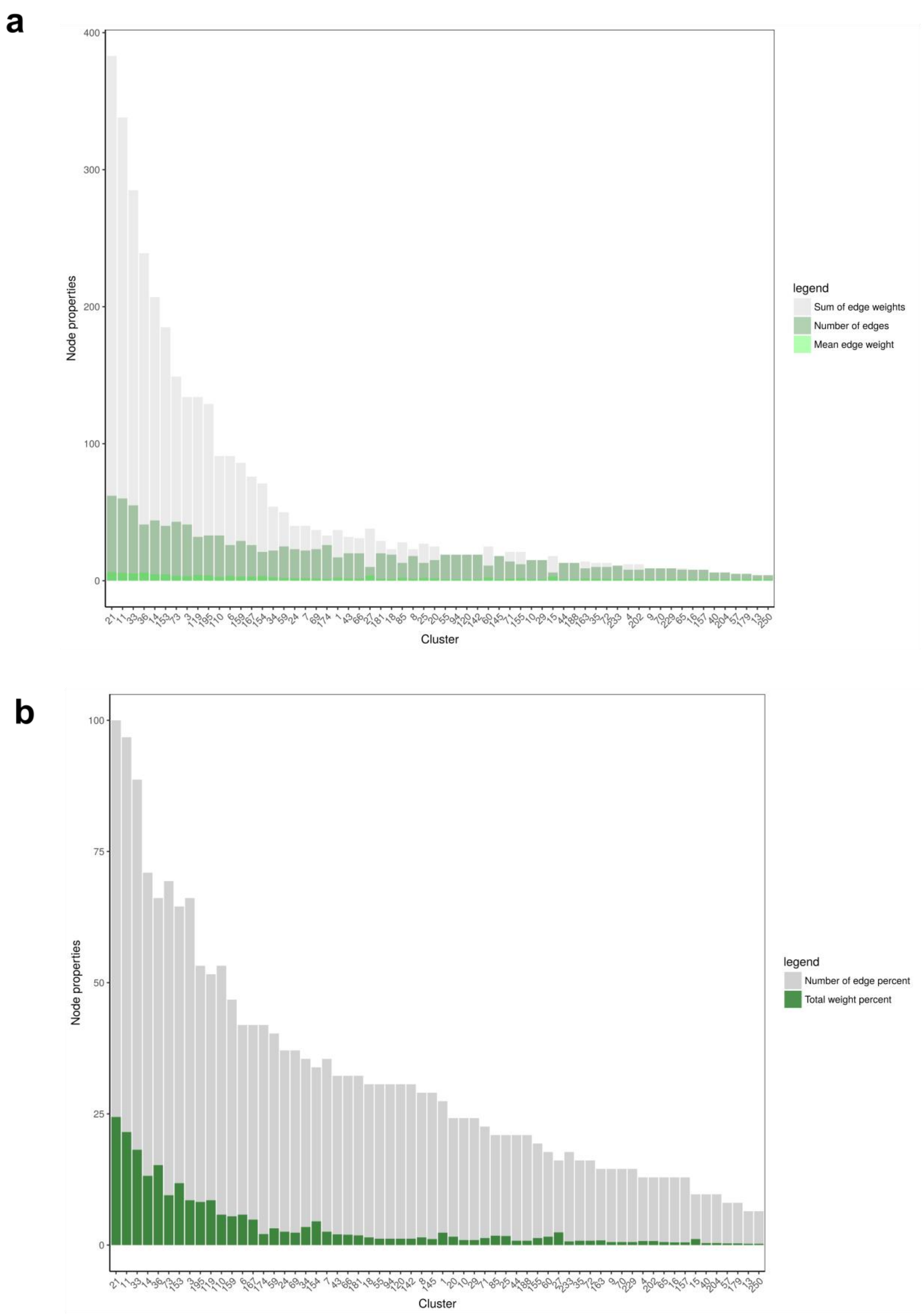


**Extended Data Figure 12.** Summary statistics of the pleiotropy graph. **a.** Total number of edges and total weight of all edges for phnotype cluster. **b.** Same data, but represented as the fraction of a total. For number of edges, numbers are realted to cluster #21 (the one with highest connectivity).

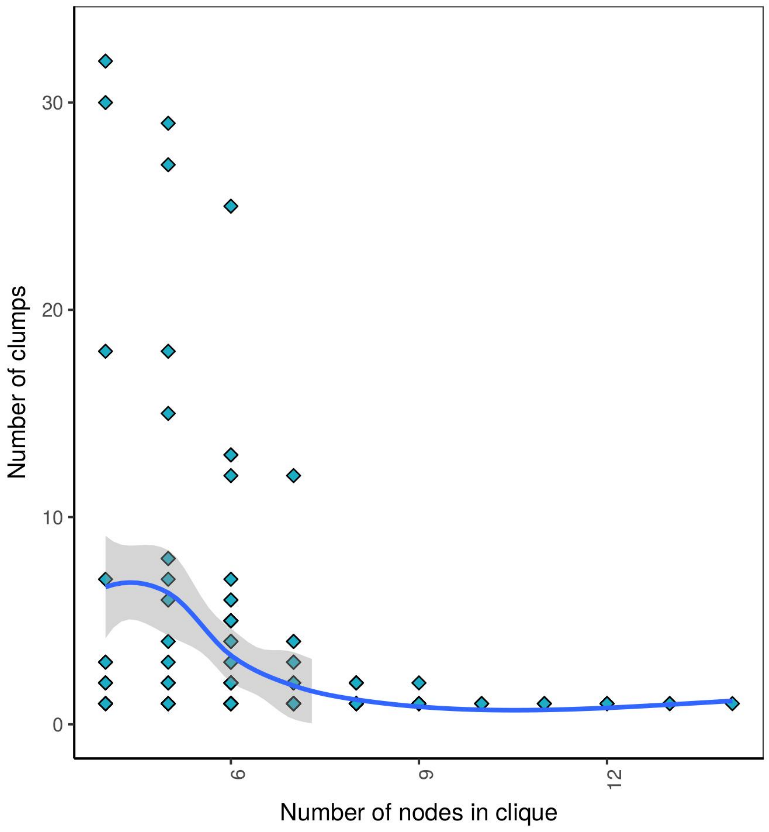


**Extended Data Figure 13.** A scatterplot of the number of shared loci depending on the number of nodes in a dense subgraph (see main text for more details). Larger groups of nodes in a network tend to share no more than 1 locus.

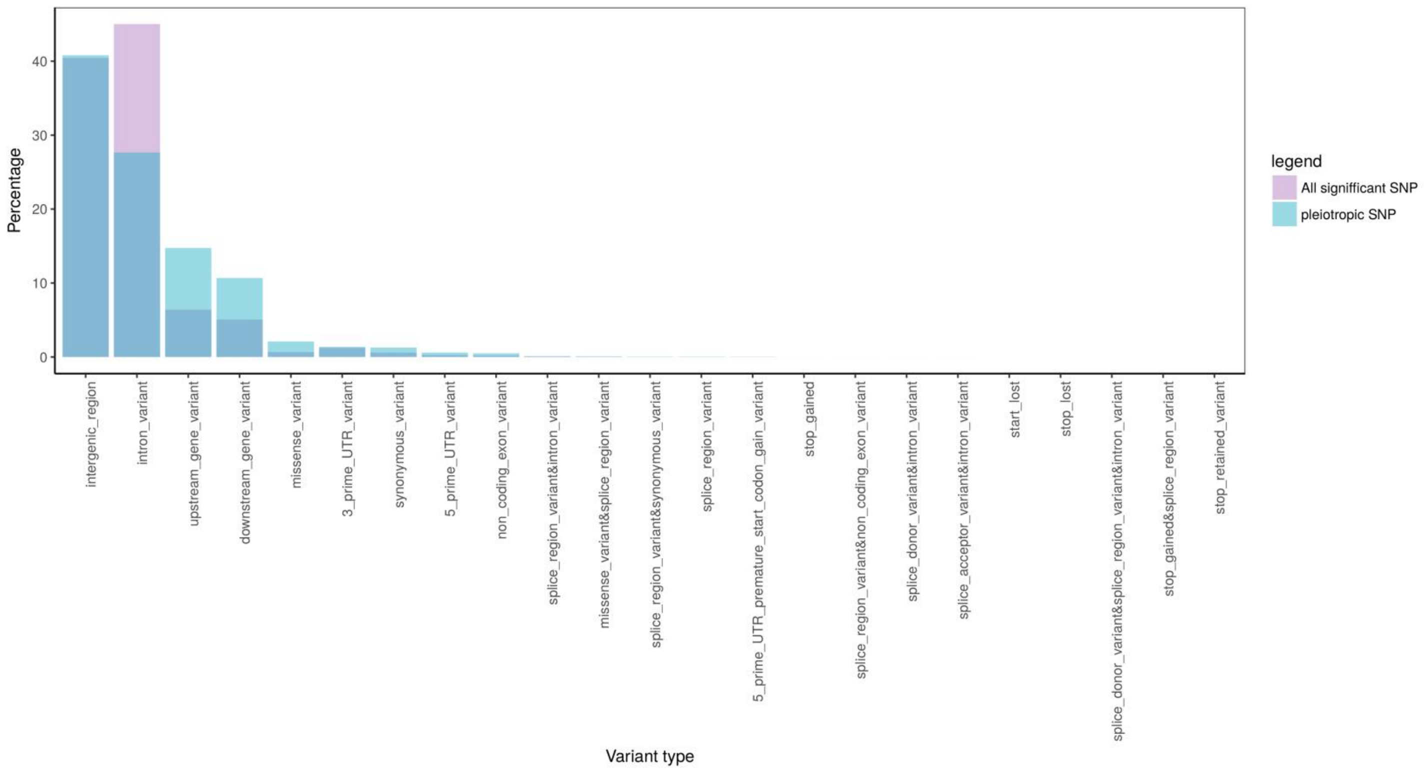


**Extended Data Figure 14.** A bar plot of the fraction of SNPs in each functional class for pleiotropic SNPs and all SNPs with at least 1 significant association. Upstream, downstream, and missense variants are overrepresented in the pleiotropic group (results are concordant with the ones repored in Watanabe et al., 2018)

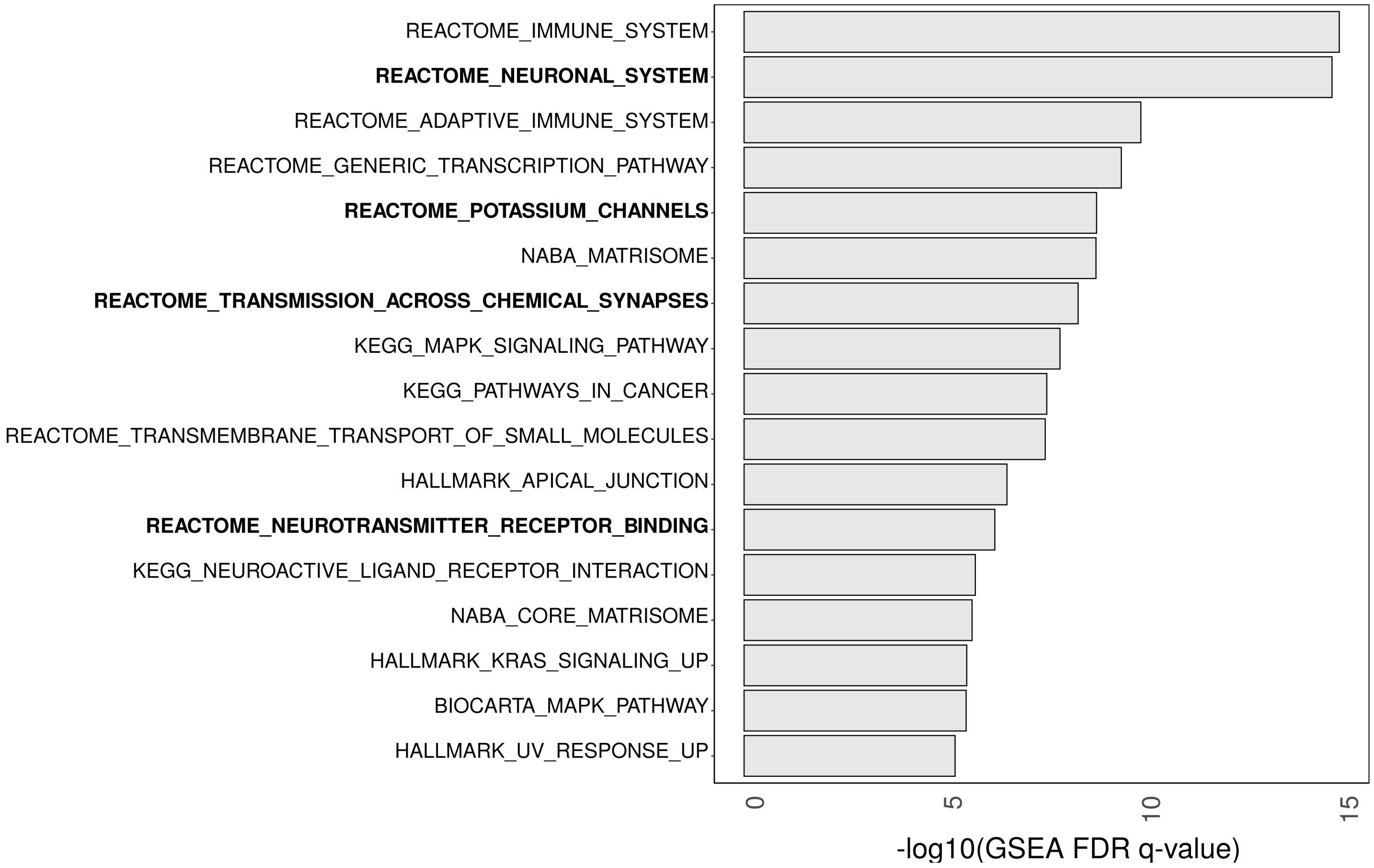


**Extended Data Figure 15.** Results of the Molecular Signatures Database (MSigDB) GSEA analysis of top-2,000 targets for the *MIR2113* microRNA (as reported in the miRBase TargetScan results). Top 10 enriched gene sets are shown. Gene sets relevant to nervous system function are highlighted in bold.

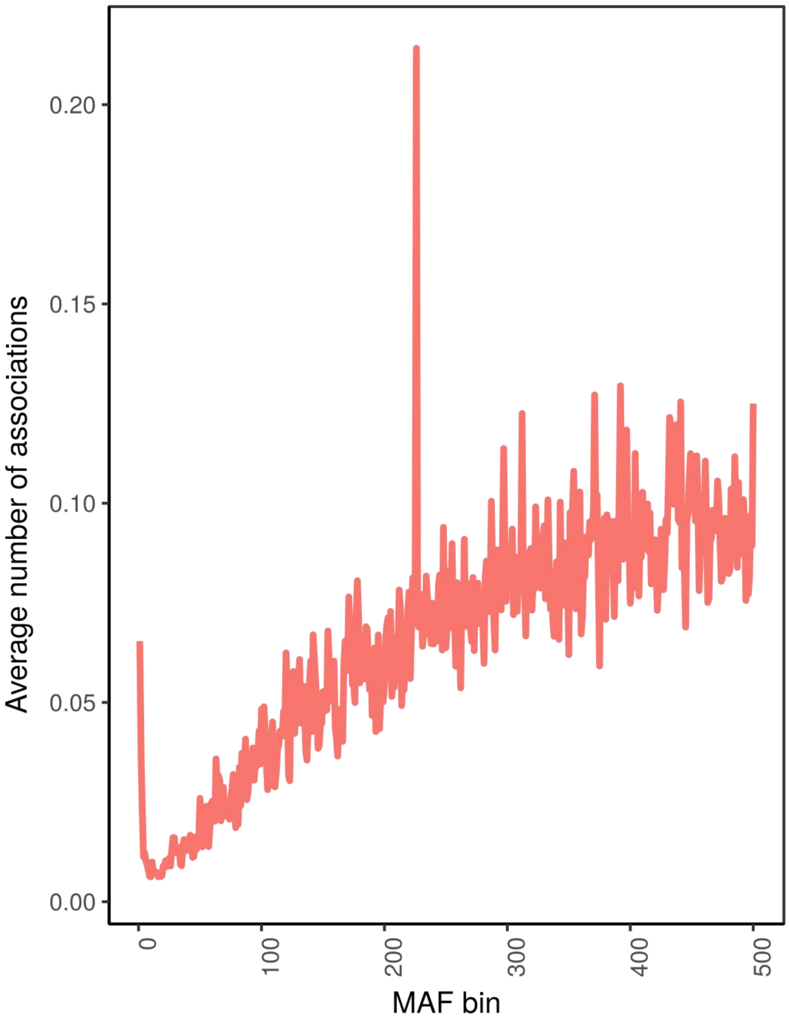


**Extended Data Figure 16.** Average number of associated clusters dependent on the minor allele frequency. MAFs were divided into 500 uniformly distributed bins ranging from 0 to 0.5.
